# Evolutionary footprints of a cold relic in a rapidly warming world

**DOI:** 10.1101/2021.07.11.451959

**Authors:** Eva M. Wolf, Emmanuel Gaquerel, Mathias Scharmann, Levi Yant, Marcus A. Koch

## Abstract

With accelerating global warming, understanding the evolutionary dynamics of plant adaptation to environmental change is increasingly urgent. Here we reveal the enigmatic history of the genus *Cochlearia* (Brassicaceae), a Pleistocene relic that originated from a drought-adapted Mediterranean sister genus during the Miocene. *Cochlearia* rapidly diversified and adapted to circum-Arctic regions and other cold-characterized habitat types during the Pleistocene. This rapid change in ecological preferences was accompanied by a highly complex, reticulate polyploid evolution, which was apparently triggered by the impact of repeated Pleistocene glaciation cycles. Our results illustrate that two early diversified arctic-alpine diploid gene pools contributed differently to the evolution of this young polyploid genus now captured in a cold-adapted niche. Metabolomics revealed ancestral central carbon metabolism responses to cold in diverse ecotypes, likely due to continuous connections to cold habitats that we hypothesize facilitated widespread parallel adaptation to alpine and subalpine habitats, and which we speculate were coopted from existing drought adaptations. Given the growing scientific interest in adaptive evolution of temperature-related traits, our results provide much-needed taxonomic and phylogenomic resolution of a model system as well as first insights into the origins of its adaptation to cold.

## Introduction

Vast spatiotemporal variation across natural environments subjects all organisms to abiotic stressors. Dynamic shifts in these stressors lead to migration, adaptation, or extinction. Thus, the current acceleration of global warming and climate volatility demands a better understanding of evolutionary dynamics resulting from climate change (Root et al. 2003; Thomas et al. 2004; Jump and Penuelas 2005; Visser 2008; Franks et al. 2014). Further, there is a strong economic rationale for understanding the consequences of environmental change on plants, which typically lack the option of rapidly migrating away from changing conditions (Xoconostle-Cazares et al. 2010; Olsen and Wendel 2013). An especially powerful natural laboratory for the study of climate change adaptation is represented by the recurrent cycles of glaciation and deglaciation during the Pleistocene. Thus, looking backwards in time by investigating the evolutionary footprints of this epoch can provide valuable insight for our understanding of adaptive evolution.

The genus *Cochlearia* L. represents a promising study system for the evolutionary genomics of adaptation not only because of a proximity to *Arabidopsis* and other Brassicaceae models, but also because of distinctive ecotypic traits which evolved within a short time span (Koch et al. 1996; Koch et al. 1998; Koch et al. 1999; Koch 2012). Among these are adaptations to extreme bedrock types (dolomite versus siliceous), heavy metal-rich soils, diverse salt habitats, high alpine regions, and life cycle variation. This diversity is accompanied by a remarkably dynamic cytogenetic evolution within the genus. Two base chromosome numbers exist (n=6 and n=7) and out of the 20 accepted taxa, two thirds are neopolyploids, ranging from tetraploids to octoploids (previous phylogenetic hypotheses are given in Supplementary Figure 1; see Supplementary Table 1 and Supplementary Note 1 for details). The connecting element between the various cytotypes and ecotypes is the cold character of the diverse habitat types standing in sharp contrast to the preferences of the sister genus *Ionopsidium*, which occurs only in arid Mediterranean habitats (Koch, 2012).

While on species-level it has been shown in *Arabidopsis thaliana* that drought- and temperature-adaptive genetic variants are shared among Mediterranean and Nordic regions (Exposito-Alonso et al. 2018) the separation of *Cochlearia* from *Ionopsidium* dates back to the mid-Miocene (Koch 2012), and the formation of the genus *Cochlearia* as we see it today first started during the middle (0.77 mya – 0.13 mya) and late (0.13 mya – 0.012 mya) Pleistocene (Koch 2012) raising the hypothesis of a long-lasting footprint of drought adaptation. The strong association with cold habitats shown by almost all *Cochlearia* species may therefore be interpreted as a cold preference that was acquired rapidly in adaptation to the intense climatic fluctuations which characterized this epoch. Such rapid response and adaptation to environmental change is of general interest, because it might be less constrained by demographic processes (Buffalo and Coop 2019).

According to the most recent taxonomic treatment (as listed in BrassiBase; Kiefer et al. 2014), the genus *Cochlearia* comprises 16 accepted species and 4 subspecies (Supplementary Table 1). Today, living in a warming world, almost all of these taxa are endangered since they strongly depend on the existence of cold habitats, which are among the first to be influenced by globally rising temperatures and the resultant transformations in abiotic conditions (Walther et al. 2002; Parmesan 2006; Alsos et al. 2012; Barrett et al. 2015).

There is a growing interest in the genus *Cochlearia* from diverse fields (Reeves 1988; Brock et al. 2006; Dauvergne et al. 2006; Brandrud et al. 2017; Mandakova et al. 2017; Nawaz et al. 2017; Bray et al. 2020), but the evolutionary history of the genus has been highly recalcitrant. Thus it is still unknown how the genus managed the rapid transition from Mediterranean to circum-Arctic or high-alpine habitat types in combination with a highly dynamic cytogenetic evolution. Here we overcome the first obstacle, presenting the first genus-wide picture, using comprehensive cytogenetic data and highly resolving phylogenomic analyses, complemented by insights into the *Cochlearia* metabolome response to cold. Our analysis uncovers a recurrent boosting of speciation by glaciation cycles in this cytotypically very diverse genus and indicates that despite clear challenges brought by global warming, the genus survives evolutionarily while rescuing its genetic diversity with reticulate and polyploid gene pools.

## Results

### Cytogenetic analyses show geographic structuring and parallel evolutionary trends towards shrinking haploid genomes in higher polyploids

In order to resolve its cytogenetic evolution, we first generated a comprehensive survey of 575 georeferenced chromosome counts and/or genome sizes across the *Cochlearia* genus (Supplementary Data Set 1) based on our novel cytogenetic data (Supplementary Table 2) and a review of published literature over the last century (Supplementary Table 3). This survey revealed a clear continental-scale geographical partitioning of diploid cytotypes (2n=12 and 2n=14; Figure 1a).

**Figure 1.**
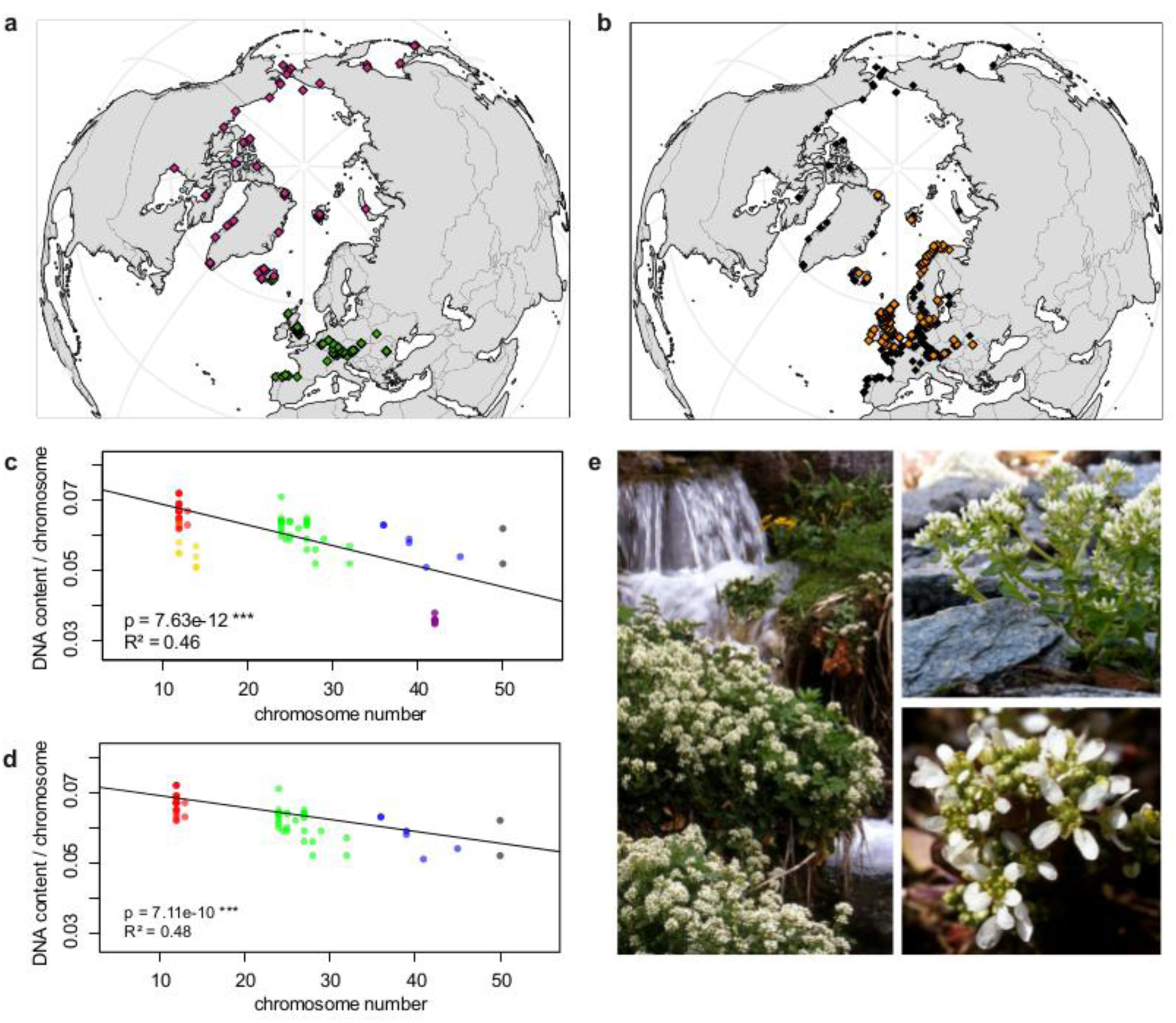
Distribution and cytogenomic flexibility of *Cochlearia*. **a**, geographic distribution of chromosome counts for diploid *Cochlearia* accessions (n=169; Supplementary Data Set 1), showing a clear separation of 2n=12 (European, green) and 2n=14 (arctic, pink). **b**, geographic distribution of aneuploidies (orange, n=138) and euploidies (black, n=376) in diploid and polyploid *Cochlearia* (n=514). **c**, linear regression analysis of measured DNA content per chromosome (given in picograms) relative to respective total chromosome numbers [red: 2n=2x (non-arctic), yellow: 2n=2x (arctic), green: 2n=4x, blue 2n=6x (excluding *C. danica*), purple: 2n=6x (*C. danica*), dark grey: 2n=8x] showing a significant decline of genome size per chromosome with increasing total chromosome numbers (78 individuals from 38 accessions representing 14 taxa analyzed in total). **d**, linear regression analysis excluding arctic diploids (yellow) and *C. danica* (purple) as putative outliers (59 individuals, 29 accessions, 11 taxa analyzed in total), showing a significant decline with increasing chromosome numbers. **e**, images of three *Cochlearia* species (left: *C. pyrenaica* (2n=2x=12), top right: *C. tatrae* (2n=6x=42), bottom right: *C. anglica* (2n=8x=48)). Figure 1―Source Data 1 This file contains the results for the distribution of cytotypes (a, b), DNA content per chromosome (c, d).

Aneuploidies (aberrations from typical species-specific chromosome numbers) are frequently found in polyploid *Cochlearia* taxa, especially along the coasts (Figure 1b), but they are only rarely spotted in diploids and therefore are nearly absent from arctic regions, where only diploids are observed (Supplementary Figure 2).

In order to further investigate the cytogenetic dynamics within *Cochlearia*, we analyzed the relationships of 1) chromosome number vs. genome size and 2) chromosome number vs. DNA content per chromosome via linear regression analyses and rank correlation tests (Supplementary Table 4). Both analyses revealed that 1) the high frequency of polyploidization events is accompanied by increasing genome sizes (Supplementary Figure 3), while there is 2) a slight but significant negative correlation of chromosome size with increasing chromosome numbers (Figure 1 c, d). This trend was independent of inclusion or exclusion of the annual species *C. danica* and the short-lived arctic diploids. These taxa were treated as putative outliers because a relationship between lower genome size and annuality was shown for the Brassicaceae as a whole (Hohmann et al. 2015).

### Organellar phylogenies provide evidence of recurrent glacial speciation boosting

To provide a highly resolved organellar phylogeny, we generated genome resequencing data for 65 *Cochlearia* accessions, representing all accepted *Cochlearia* species, as well as three species from the sister genus *Ionopsidium* (Supplementary Figure 4 and Supplementary Data Set 2). Complete chloroplast genomes were assembled *de novo* for all samples. A maximum likelihood (ML) analysis using RAxML based on the whole chloroplast genome alignment (122,798 bp, excluding one copy of the inverted repeat) covering a total of 5,292 SNPs (1,003 within *Cochlearia*) revealed six well-supported major lineages within the genus (Supplementary Figure 5, congruent to lineages as illustrated in BEAST chronogram, Figure 2). A rapid radiation was indicated by the existence of several polytomies, despite the generally high resolution of the ML tree. To test if the phylogenetic scenario as revealed by chloroplast genome analyses was also supported by the mitochondrial genome, we generated a mitochondrial ML phylogeny based on a combination of *de novo* assembly and referenced-based mapping (Supplementary Figure 6). The two maternal phylogenies are largely consistent, and after collapsing all branches below a bootstrap support of 95%, no incongruences remained (illustrated by a tanglegram in Supplementary Figure 7).

**Figure 2.**
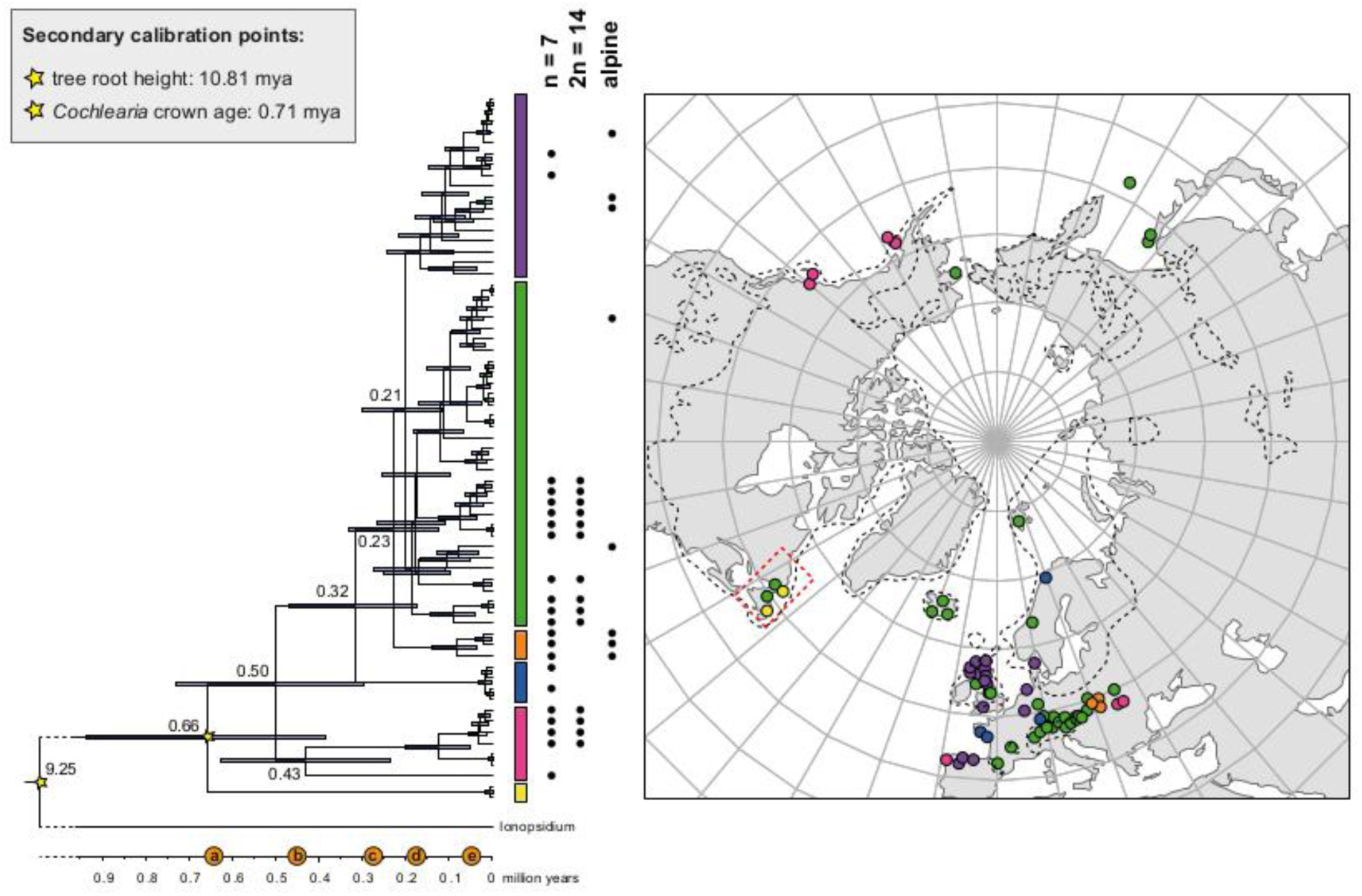
The maternal footprint of recurrent glacial speciation boosting. BEAST chronogram based on complete plastid genome sequence data supplemented by a geographical distribution pattern of the six main phylogenetic lineages (displayed as colored bars next to the tree; dots in the map are colored accordingly). The *Ionopsidium* outgroup lineage is collapsed and condensed. The tree topology is congruent to the topology as revealed from maximum likelihood (ML) analysis. The full BEAST chronogram and the ML tree (incl. bootstrap support values) are given with Supplementary figures 5, 8 and 9. Individuals with a base chromosome number of n=7 as well as diploids with 2n=14 and accessions with an alpine or subalpine habitat type are marked with black dots next to respective tips. Letters a) to e) as displayed on the timeline indicate high glacial periods: a) 640 kya, end of Günz glacial; b) 450 kya, beginning of Mindel glacial; c) 250-300 kya, Mindel-Riss inter-glacial; d) 150-200 kya, Riss glacial; e) 30-80 kya, Würm glacial. The black dashed line indicates the extent of the Last Glacial Maximum (LGM ∼ 21 kya; based on Ehlers and Gibbard (2007)). The red dashed rectangle highlights a region with evidence for ice free areas during the Penultimate Glacial Period (∼140 kya; based on Colleoni et al. (2016)).

We generated divergence time estimates with BEAST based on the whole chloroplast genome alignment. We used two secondary calibration points (*Ionopsidium*/*Cochlearia* split: 10.81 mya; *Cochlearia* crown age: 0.71 mya) taken from a large-scale age estimation analysis performed by Hohmann et al. (2015) which included 5 *Cochlearia* samples and one *Ionopsidium* sample that are also included in the present study. The revealed tree topology (Figure 2; full tree given with Supplementary Figures 8 and 9) is congruent with the topology of the ML tree. In accordance with Hohmann et al. (2015), our BEAST analysis shows a diversification of the entire genus within the last ∼660 kya, after a long period of evolutionary stasis and zero net diversification following a deep split from the genus *Ionopsidium* ∼9.25 mya (see also Koch [2012]), and in concert with the beginning of the Pleistocene’s major climatic fluctuations, which are dated to 700 kya (Webb III and Bartlein 1992; Comes and Kadereit 1998). Diversification times in the six major chloroplast lineages as revealed via ML and BEAST analysis closely coincide with high glacial periods (Petit et al. 1999; see timeline in Figure 2; Augustin et al. 2004).

The most basal *Cochlearia* chloroplast and mitochondrial haplotypes were found in Eastern Canadian *C. tridactylites*, a species of unknown ploidy (Supplementary Note 2), in a region where ice-free areas putatively occurred during the Penultimate Glacial Period (∼ 140 kya; Colleoni et al. 2016). The earliest diverged organellar genomes of known diploids were found in the arctic species *C. groenlandica* and *C. sessilifolia* (collected in British Columbia, Canada and Kodiak Island, Alaska respectively; pink lineage in Figure 2, see Supplementary Figure 5 for details), with distribution ranges covering areas such as Beringia that were thought to have served as ice-free Pleistocene refugia (Abbott and Brochmann 2003). European diploids are found in the derived green and purple lineages only. Some of the European polyploids, however, harbor early diverged haplotypes from the otherwise arctic pink lineage. Thus, except for the eastern European *C. borzaeana* with 2n=8x=48, all taxa from the pink lineage, as well as several taxa from the early diverged blue and orange lineages, have a base chromosome number of n=7 (see Figure 2).

### Genomic data and demographic modelling indicate glacial expansion

In order to analyze the nuclear fraction of our resequencing data, we mapped reads of each sample to our previously published *C. pyrenaica* transcriptome reference (total length: 58,236,171 bp; Lopez et al. 2017)). For 63 samples with sufficient nuclear sequence data quality (62 *Cochlearia* samples and *Ionopsidium megalospermum*), we generated a phylogenetic network using SplitsTree (Huson 1998; Huson and Bryant 2006) based on 447,919 biallelic SNPs. Concordant with our cytogenetic results, the network shows a clear separation of arctic and European diploid taxa (Figure 3a and b; see Supplementary Figure 10 for detailed SplitsTree output). Close associations of both *C. tridactylites* and *C. danica* with *Ionopsidium* support the picture as revealed from organellar phylogenies. Further support for the early divergence of these two species came from a maximum likelihood analysis based on 298,978 variant sites (same set of samples) performed via RAxML with an ascertainment bias correction and a general time-reversible substitution model assuming gamma distribution (Supplementary Figure 11).

**Figure 3.**
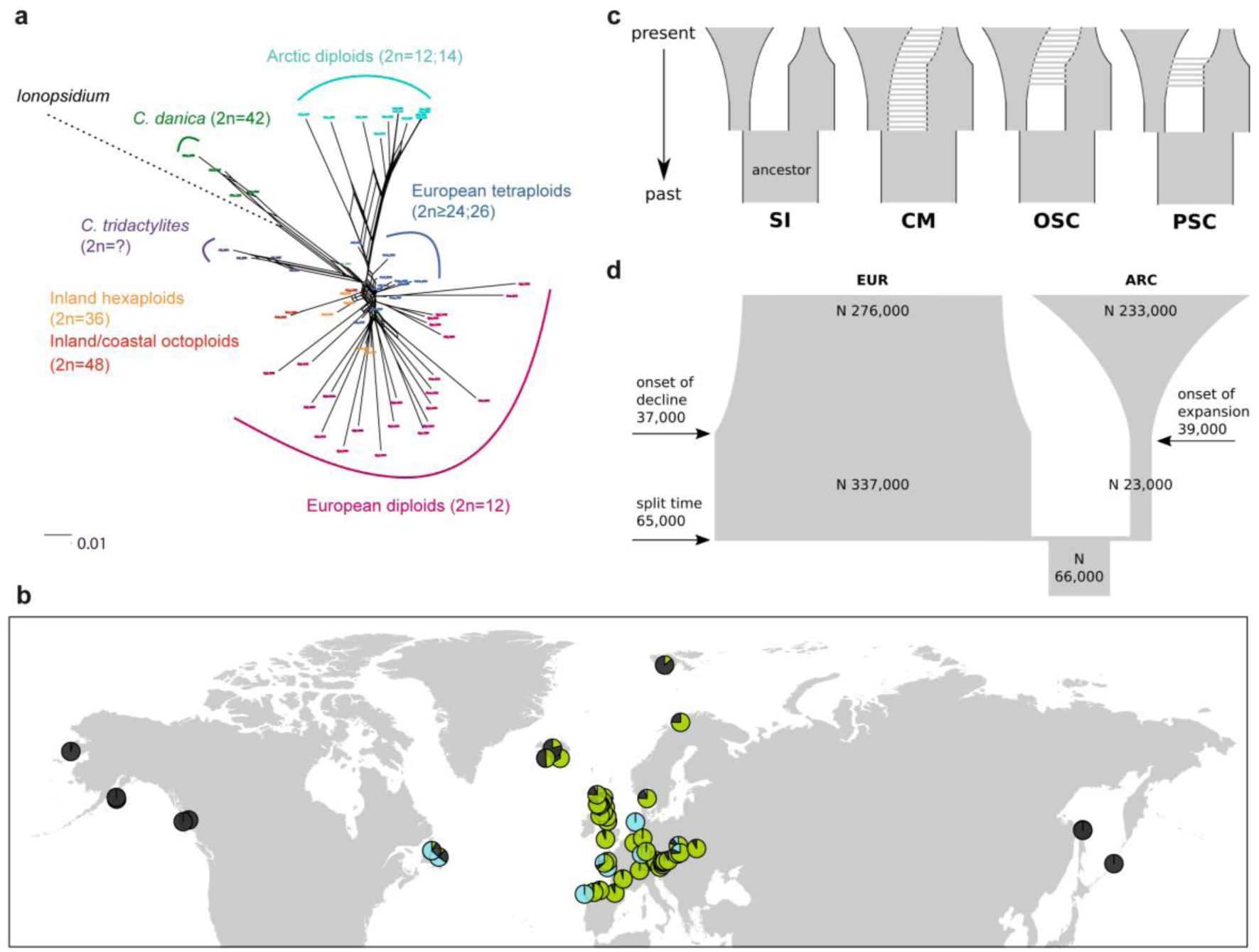
Demographic structure and history of the *Cochlearia* genus based on nuclear genome sequence data. **a,** SplitsTree analysis of 62 *Cochlearia* samples and *Ionopsidium* (outgroup) using the NeighborNet algorithm based on uncorrected p-distances (network with tip labels is given with Supplementary Figure 10). **b,** Geographic distribution of 62 *Cochlearia* samples. Chart colors correspond to STRUCTURE results (62 *Cochlearia* samples, 400,071 variants) at *K*=3; black: arctic gene pool, green: European gene pool, blue: *C. danica*-specific cluster (STRUCTURE result with tip labels given with Supplementary Figure 15). **c,** Coalescent models for diploid populations explored with Approximate Bayesian Computation (ABC). SI = strict isolation, CM = continuous migration from the population split to the present, OSC = ongoing secondary contact with gene flow starting after population split and continuing to the present, PSC = past secondary contact with gene flow starting after population split and stopping before the present. **d,** Most likely demographic history of diploid EUR (2n=12) and ARC (2n=14) *Cochlearia* populations from coalescent modelling (based on 22 EUR individuals and 12 ARC individuals; 2,140 SNPs at four-fold degenerate sites) and ABC of models without gene flow and upper bound of the population size (N) prior as 400,000 (Supplementary Table 8). N is in number of diploid individuals, and time in number of generations ago.

A STRUCTURE analysis of *Cochlearia* samples only (same variant calling, 400,071 variants after excluding *Ionopsidium*) with *K*=3 (optimal K following Evanno Method; Evanno et al. 2005) revealed a pattern very similar to that obtained via SplitsTree (Figure 3b). It shows the two diploid clusters and a third cluster comprising *C. danica* samples. Signatures of admixture between the diploid clusters can be found, especially in Icelandic 2n=12 and 2n=14 diploids and in several polyploid samples (discussed in detail in Supplementary Note 1). Interestingly, *C. tridactylites*, the earliest diverged lineage according to the plastome analysis, was modelled as a mix of the arctic gene pool and *C. danica*. In a separate TreeMix (Pickrell and Pritchard 2012) analysis of all *Cochlearia* accessions and *Ionopsidium* (447,919 variants; up to 10 migration events) the bootstrapped graph for m=1 (optimal number of migration events according to Evanno Method) likewise indicates that *C. tridactylites* is admixed, with a majority ancestry from near the base of the European (excepting hexaploid *C. danica*) and arctic groups, and a minority ancestry from the 2n=14 group of arctic diploids (see Supplementary Figures 12 to 14).

In order to elucidate the early stages of the *Cochlearia* species complex formation that might have facilitated the general cold association of the genus, we tested hypotheses regarding the evolutionary history of the diploid lineages from the arctic (ARC) and European (EUR) distribution ranges by modelling possible histories using a coalescent framework with Approximate Bayesian Computation (ABC; Tavaré et al. 1997; Beaumont et al. 2002). For a dataset of 22 European (2n=12) and 12 arctic (2n=14) individuals (Supplementary Table 5 for sampling), we analyzed 2,140 SNPs at four-fold degenerate sites (Methods). Overall, the EUR metapopulation exhibits much higher genetic diversity than the ARC metapopulation (Supplementary Table 6). In both populations we found an excess of rare alleles (negative Tajima’s D, Supplementary Table 6), indicating that they are not in mutation-drift equilibrium. Differentiation and divergence between ARC and EUR were overall very low (Fst ∼0.098, dxy ∼0.0036, Supplementary Table 6).

Given the dynamic nature of their ice-age-associated speciation histories, we hypothesized that after the EUR and ARC metapopulations separated, they underwent dramatic changes in effective population sizes (*N*_e_) over time, and that they experienced gene flow. We first tested four different hypotheses regarding the occurrence and relative timing of gene flow between ARC and EUR, because failure to account for gene flow can confound the inference of population size changes. Our gene flow hypotheses were formulated as different model categories (Figure 3c), and random forest ABC (ABC-RF; Marin and Pudlo 2015) was used to test under which model the observed data was most probable to have arisen. However, discriminating between the four gene flow models was ambiguous (see Supplementary Table 7), as the most probable model depended on the priors, particularly on the upper bound of the *N*_e_ priors. When allowing *N*_e_ up to 400,000, a model of ongoing secondary contact (OSC) prevailed over a model without any gene flow (model SI; posterior probability > 0.75, i.e. Bayes Factor > 3), but OSC was not clearly better than a model with continuous gene flow (CM) or a past secondary contact (PSC). Yet, when choosing a less informative *N*_e_ prior with a greater upper bound of three million, all four models were similarly in agreement with the observed data. In the absence of strong prior information for *N*_e_, we could not establish the occurrence of gene flow between ARC and EUR metapopulations with confidence (see Supplementary Note 3 for further explanations on ABC model choice).

To estimate changes in *N*_e_ through time, we fitted parameters of a model without any gene flow (SI) and a model of OSC, considering both high and low *N*_e_ upper prior bounds, amounting to a total of four model fits (Figure 3c; see Supplementary Table 8). The general pattern of changes in *N*_e_ through time were always modelled such that each of ARC and EUR had an older phase of constant *N*_e_ followed by a recent phase of exponential expansion or decline.

The model without gene flow (SI) contained the fewest parameters, and this model with a smaller *N*_e_ prior bound of maximal 400,000 provided the best fit to the data (smallest Euclidean distances between observed and predicted values from posterior predictive checks; see Supplementary Table 9). This model fit (Figure 3d) is consistent with the remaining three model fits. Importantly, all model fits agreed about the relative *N*_e_ of EUR and ARC: they evolved drastically differently, with the EUR metapopulation having risen to 4 to 12-fold the *N*_e_ of their common ancestral population followed by a moderate decline (0.4 to 0.8-fold in three out of four models), or constant size up to the present (OSC with smaller *N*_e_ upper bound). In contrast, the ARC metapopulation experienced a bottleneck after splitting from the common ancestor (0.2 to 0.5-fold), followed by a dramatic expansion of 9- to 52-fold (Supplementary Table 8). Estimated *N*_e_ for the ancestral population and for ARC during the ancient phase were robust to the choice of model and priors, but other *N*_e_ parameters, in particular the ancient phase of EUR and the recent phase of ARC (i.e. the phases in which their *N*_e_ were largest), increased when the prior’s upper bound was increased. However, the relative *N*_e_ trends through time were robust to these uncertainties, as mentioned above.

If EUR and ARC did not experience gene flow (SI model fit), they must have separated only about 65,000 to 73,000 generations ago, corresponding to 0.2 to 3 *N*_e_ units. This estimate was robust to the choice of priors and may coincide with the last interglacial period (considering a 2 year average generation time; Abs 1999). If gene flow is assumed (OSC), this split could have occurred earlier (119,000 to 227,000 generations ago; Supplementary Table 8). Further parameters were poorly estimable as indicated by large prediction error, and little deviation between prior and posterior. These include the timing of the transitions from ancient to recent phases of *N*, the timing of migration, and the migration rates. Considering our BEAST analyses from plastome data, a split-time of 65,000 to 73,000 generations ago is more likely (e.g. dating of splits within the green evolutionary lineage). An important implication of this is that the polyploid inland taxa mediating between the two diploid gene pools did not evolve earlier than during the last Interglacial.

### Genus-wide cold response characterized by metabolic profiling indicates an ancient origin of cold temperature tolerance

We hypothesized that a very early evolved tolerance to cold facilitated the observed widespread parallel adaptation to alpine and subalpine habitats across the *Cochlearia* genus. To test this hypothesis, we performed metabolite profiling of central carbon metabolism using gas chromatography-mass spectrometry (GC-MS) for 28 worldwide *Cochlearia* accessions, including 14 taxa and five outgroup accessions (genus *Ionopsidium;* sampling details in Supplementary Table 9). For the *Cochlearia* accessions, we defined four climatic ecotypes based on a hierarchical cluster analysis of 9 WorldClim bioclimatic variables related to temperature and temperature/precipitation (Figure 4a and b; see Source Data for Figure 4a). *Ionopsidium* represented a fifth, Mediterranean, ecotype.

**Figure 4.**
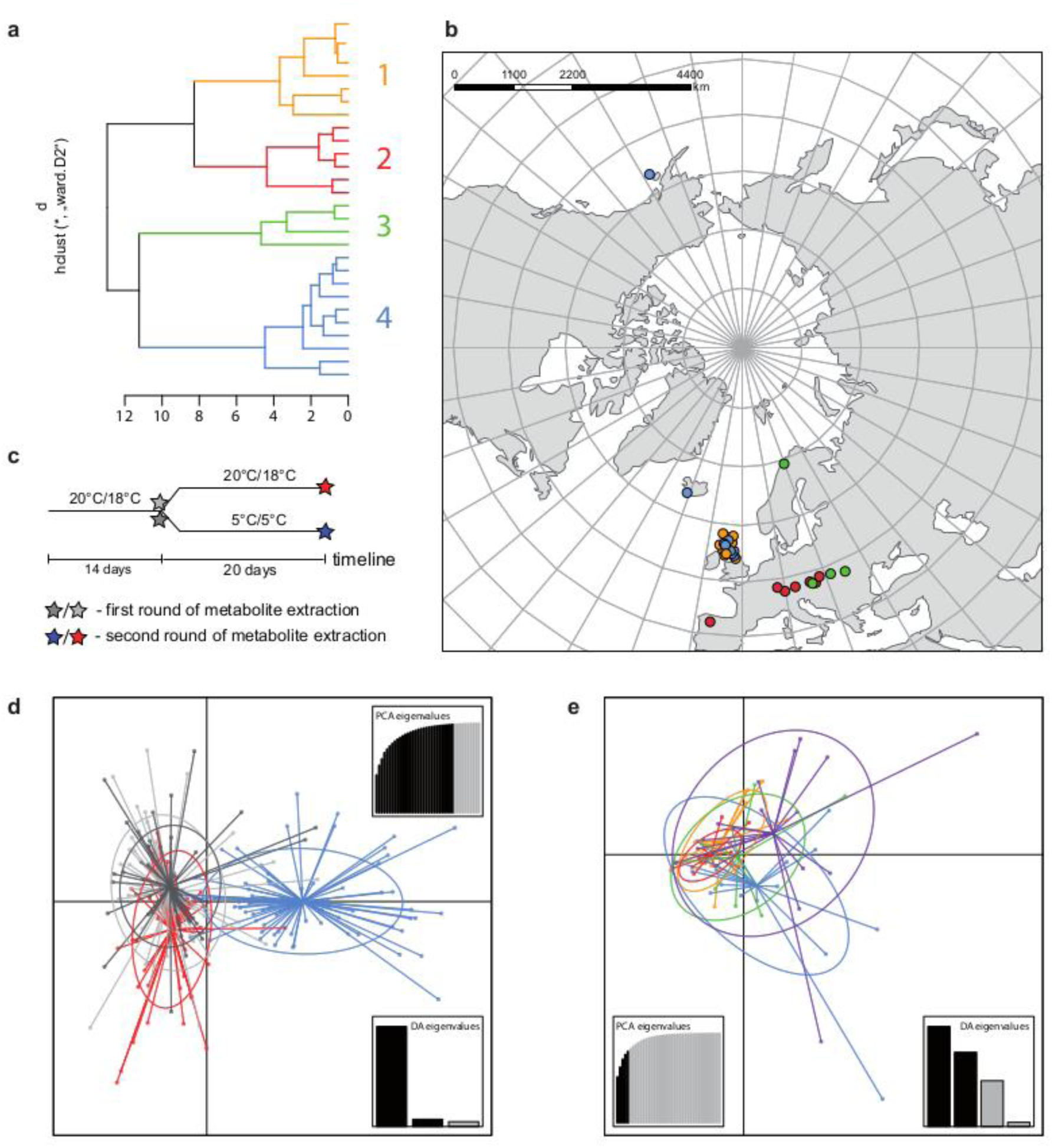
A common cold response indicated by metabolic profile clustering. **a,** hierarchical cluster analysis of 28 *Cochlearia* accessions based on 5 temperature-related and 4 temperature/precipitation-related bioclimatic variables (WorldClim; given with Source Data) using Euclidian distances and the Ward’s method (Ward Jr 1963; Murtagh and Legendre 2014). Cluster 1: coastal accessions of polyploids from the northern UK; Cluster 2: European inland accessions (diploids and polyploids); Cluster 3: European accessions with arctic/alpine habitat types (Norway, Carpathians, High Tatra Mountains, Austrian Alps); Cluster 4: arctic accessions (Iceland, Alaska) and alpine habitat types from the UK. **b**, geographical distribution of bioclimatic clusters as discovered via hierarchical cluster analysis (colors are representing clusters 1-4, see **a**). **c**, experimental setup of temperature treatment with a timeline for metabolite extractions. **d**, DAPC based on all metabolic profiling measurements, grouped by treatment (colors are representing metabolite extractions as illustrated in c). **e**, DAPC of metabolite measurements after cold treatment grouped by 4 bioclimatic clusters (colors as in **a)** and **b)**) and *Ionopsidium* (purple). Figure 4―Source Data 4 This file contains the bioclim data used for cluster analysis (a), and metabolic profiling data (d,e) used for DAPC analyses.

Metabolic profiles were collected before and after a 20 day treatment under cold (5°C) or control conditions (18°C/20°C; Figure 4c). From these profiles we exported peak areas for a set of 40 compounds consistently detected across our samples that were annotated as intermediates within central carbon metabolic pathways (see Supplementary Table 10 for a list of detected compounds; raw/normalized data provided in Supplementary Data Set 3).

As has been best seen in the *Arabidopsis* cold stress metabolome (Cook et al. 2004; Kaplan et al. 2004; Kaplan et al. 2007), we found increases in the relative levels of many of the detected compounds after the cold treatment, especially for carbohydrates and amino acids (data grouped by bioclimatic clusters, analyzed by one-way ANOVAs; Supplementary Figures 16 to 21).

Initially, we ran PCA analyses with different group priors for the 40 compound matrix, yet for most of the pre-defined clusters these analyses revealed little discriminating structure (see Supplementary Figures 22 and 23). Thus, to increase power, we performed discriminant analyses of principal components (DAPC), thereby maximizing the variation between clusters while minimizing the variation within (Jombart et al. 2010). When grouping the data by temperature treatment, DAPC shows a strong metabolomic response of all clusters to temperature stress (cold treatment at 5°C; Figure 4d; Supplementary Figure 24 and Source Data). Yet, surprisingly, no demarcation of the cold responses could be revealed either among the four *Cochlearia* ecotypes or even between *Cochlearia* and *Ionopsidium* when data were grouped according to bioclimatic clusters (Figure 4e). Because both drought, salt and cold stress strongly involve osmotic challenge these data support the idea that an evolution to one of these stressors in the common ancestor not only allowed for the survival under Mediterranean conditions, but also may have preadapted the nascent *Cochlearia* genus to habitats characterized by cold or salt.

## Discussion

Previous studies provided first insights into the complex evolutionary history of the genus *Cochlearia* (e.g. Koch et al. 1996; Koch et al. 1998; Koch et al. 1999; Koch 2002; Koch et al. 2003; Kochjarová et al. 2006; Cieślak et al. 2007; Koch 2012; Brandrud et al. 2017; Bray et al. 2020). Yet, a genus-wide picture has been missing. Here we incorporate cytogenetics, phylogenomics and metabolomics to present the complex and dynamic evolutionary past of this cold adapted, rapidly radiating genus.

As revealed from the continental-scale geographical partitioning of diploid cytotypes (2n=12 and 2n=14; Figure 1a) we see an early split of two diploid gene pools followed by separation of the two diploid metapopulations in arctic regions (2n=14) and in Europe (2n=12). Only in Iceland do both diploid cytotypes co-occur. Both cytotypes underwent genome and chromosome size reduction within this arctic environment, which may be seen as a metabolic advantage (e.g., meiotic cell division under cold stress (Bomblies et al. 2015), or triggered by shorter life cycles). The exclusively circum-Arctic 2n=14 taxa may have also benefitted from one extra centromere, which increases total recombination rate and thus may also be favorable under harsh conditions and drastic environmental changes (Stapley et al. 2017). As also shown below from a phylogenetic point of view, the most parsimonious scenario recognizes 2n=14 and thus a base chromosome number of n=7 as ancestral state (Figure 2).

Whereas the presence of chromosomal aberrations in *Cochlearia* have been postulated to be caused by B-chromosomes (Gill 1971b; 1973; Gill et al. 1978; Nordal 1988), these aneuploidies could also be the result of hybridization/polyploidization events followed by mitotic/meiotic difficulties or independent ascending/descending dysploidies via chromosome rearrangements (Lysak 2014). Karyotypic variation and aneuploidy following hybridization in plants have been described as adjustment processes within the allopolyploid genome putatively connected to the beginning of re-diploidization (Mandáková et al. 2016). For *Cochlearia*, the high frequency of chromosomal aberrations in polyploids likely reflects a highly dynamic genome evolution, given the near absence of interspecific fertility barriers between cytotypes (Gill 1971a; 1973; Koch et al. 1998). Moreover, a trend for reducing genomic redundancy in the young polyploids is indicated by the negative correlation of chromosome size with increasing chromosome numbers. Combined with the stable diploid karyotypes, these patterns are well in accordance with the hypothesis of a necessary balance between both genomic flexibility and stability allowing for further adaptive evolution to a continuously changing environment (Hohmann et al. 2015).

The observed tight concurrence of high glacial periods and diversification times within the six major lineages (Figure 2) suggests a boosting impact of these climatic fluctuations on the diversification of the genus. Accordingly, the observed polytomies, especially within the green and the purple lineage, likely reflect events of rapid, reticulate radiation during the last two glacial periods, where we speculate that polyploidization and hybridization may have facilitated adaptation to dramatically changing environments (Hohmann et al. 2015).

Based on the organellar phylogenies and divergence-time estimates (and contradicting former hypotheses (Gill 1971a; 1973; Koch et al. 1996)), an evolutionary scenario with an ancestral base chromosome number of n=7 in diploid arctic ancestors followed by reduction to n=6 in European diploids is most parsimonious. This view is also supported by the finding that also within the green lineage, basal groups are confined to arctic regions. However, in accordance with former hypotheses (Koch 2002; Koch et al. 2003), we can also postulate that additional glacial refugia must have existed in Europe north of the primary refugia for European species (Iberian Peninsula, Southern Italy, Balkans; Taberlet et al. 1998)), as also described for other taxa (Stewart and Lister 2001; Tribsch et al. 2002; Tribsch and Schönswetter 2003; Provan and Bennett 2008). The putative arctic background of the entire genus *Cochlearia* might have a) promoted the general association with cold-characterized habitat types throughout the genus and b) facilitated later adaptation to different high alpine regions in Europe. As revealed from the plastid phylogeny (Supplementary Figure 5), the latter has taken place repeatedly in the High Tatra Mountains (*C. tatrae*, orange lineage), in the Austrian Alps (*C. excelsa*, green lineage), in alpine habitat types in Great Britain (*C. alpina*/*C. micacea*, purple lineage), and in several other European habitats with a more subalpine character, namely in the Pyrenees or in Switzerland.

With respect to the putative arctic origin of the genus in its current shape with an ancestral base chromosome number of n=7, the very strong admixture between both diploid genetic clusters of different chromosome numbers (as seen in diploid arctic Icelandic accessions) provides evidence of genetic coherence. Considering the numerous polyploids and their independent, multiple origins in *Cochlearia* (see Supplementary Note 1) and the exceptional case of sympatry of both different diploid cytotypes in an arctic environment on Iceland, it seems that the two early separated gene pools serve as a continuous source for the evolution of new, often transient, and currently highly endangered species.

Regarding the demographic histories of the two gene pools, our ABC data suggest that ARC and EUR metapopulations are only weakly differentiated and share most of their genetic variation, either because of recent separation or gene flow. They currently have large and similar effective population sizes, but ARC has expanded after a bottleneck whereas EUR has remained constant, or recently declined.

When exposed to low but non-freezing temperatures, plants commonly activate cold acclimation responses involving changes in gene expression, enzymatic activities and ultimately, concentrations of many central carbon intermediates (Kaplan et al. 2004; Kaplan et al. 2007; Hoermiller et al. 2017). As part of this low temperature response, particular accumulations of carbohydrates, polyols and amino acids are detected (Kaplan et al. 2004; Krasensky and Jonak 2012).

Given that all *Cochlearia* ecotypes, regardless of origin, exhibited similar responses to the temperature treatment performed under controlled conditions for central carbon intermediates, we hypothesize that these similar responses to cold within the genus reflect continuous connections to cold-characterized habitats acquired over the course of a migration to northern/circum-Arctic regions during the early evolution of the genus. Apparently, a constitutive cold tolerance has not been lost secondarily within *Cochlearia*. Future analyses of the cold-induced metabolome of *Cochlearia* will require untargeted metabolomics analyses to systematically account for the contribution of species/ecotype-specific metabolic characters in the overall cold response. In this respect, previous studies on glucosinolate and tropane alkaloid chemotypes, which are key to biotic stress adaptation, have shown rapid structural diversification and ecotype specificities within the *Cochlearia* genus (Brock et al. 2006; Blažević et al. 2020). Regarding the overall similar response of central carbon metabolism to the cold treatment of annual species from the genus *Ionopsidium* which are adapted to at least seasonally dry habitats and/or salt habitats, we may conclude that commonalities between cold and salinity/drought stress would enable a putative recruitment of the cold tolerance from existing adaptations to drought and/or salt as hypothesized for other plant taxa (Böndel et al. 2018; Exposito-Alonso et al. 2018).

## Conclusion

Our results indicate that the immense karyotypic diversity of *Cochlearia* reflects not only recurrent hybridization, but also speciation boosting during Pleistocene glaciation cycles. This in turn triggered parallel exploration of new ecological niches on a continental scale ranging from the Arctic towards lower latitude alpine regions. Genetic diversity was meanwhile rescued in reticulate and polyploid gene pools, often tearing down species boundaries. Irrespective of their habitat contrasts, both *Cochlearia* and its Mediterranean sister genus *Ionopsidium* show a pronounced response to cold stress based on metabolite profiling. This leads to the idea that a shared ancestral adaptation to drought enabled not only the survival under dry Mediterranean conditions, but also may have preadapted the nascent *Cochlearia* genus to habitats characterized by cold or salt: all of these stressors have at their cores osmotic challenge. As colonization of these habitat types occurred multiple times independently, *Cochlearia* represents a powerful system to predict the fate of this and other taxa in a world marked by climate change.

## Methods

### Plant Material and Taxon Sampling for Cytogenetic Analyses

Our cytogenetic analyses (chromosome counts and flow cytometry) were performed on a large collection of living plants including different ploidy levels and covering big parts of the genus’ distribution range. The living plants, either collected in the wild or grown from seeds in the Botanical Garden Heidelberg, were cultivated under greenhouse conditions in a substrate composed of seedling potting soil, quartz sand, and either composted earth or Ökohum’s peat-based substrate (Ökohum GmbH, Herbertingen, Germany). During the summer season, we transferred parts of the plant collection to a plant room with a controlled 16/8h day/night cycle under 20°C.

Aside from own data, we performed a literature survey for published chromosome counts/genome size measurements considering literature from the last 100 years (Supplementary Table 3). Published and own datapoints were finally merged in a comprehensive database (Supplementary Data Set 1) and if possible, coordinates of the respective population localities were added as listed in the literature or carefully approximated based thereon. Every population listed by one author was treated as an individual datapoint. This way, we collected 575 georeferenced database entries and visualized these using the R package ‘ggplot2’ (Wickham 2016; R version 3.3.1).

### Flow Cytometry

In order to minimize the risk of contaminating factors such as fungi and to reduce the generally high amount of endopolyploidy, we selected only very young and healthy leaves for flow cytometry sample preparation. Nuclei extraction and staining were performed on ice using the Partec two-step kit CyStain PI Absolute P (Partec GmbH, Münster, Germany) following manufacturer’s instructions with minor modifications. Namely, 15 mM β-mercaptoethanol and 1% w/v polyvinylpyrrolidone 25 (PVP) were added to the staining buffer. After chopping leaf material of each sample together with the chosen internal standard plant in 500 μl of the extraction buffer, the suspensions were filtered through 50 μm CellTrics filter (Sysmex Partec GmbH, Görlitz, Germany). 2000 μl of the staining buffer were added to each flow-through and samples were incubated on ice for 30 to 60 minutes.

A list of standard plants used for flow cytometry experiments together with respective 2C-values is given with Supplementary Table 11. In order to ensure consistent measurements, 2C-values of all standard plants were finally readjusted to *Solanum*, which was used as a reference.

For several samples, instead of using the Partec kit, an alternative lysis buffer LB01 as specified by Doležel et al. (1989) was used, containing 15 mM Tris, 2 mM Na2EDTA, 0.5 mM spermine tetrahydrochloride, 80 mM KCl, 20 mM NaCl, 0.1% v/v Triton X-100; 15 mM β-mercaptoethanol and 0.1 mg/ml RNAse A added right before preparation. Both buffers resulted in similar peak qualities, generated in a Partec CyFlow Space flow cytometer (Sysmex Partec GmbH) using a 30 mW green solid state laser (λ=532 nm). Gating and peak analysis were performed using the Partec FloMax software version 2.4 (as exemplified in Supplementary Figure 25; detailed flow cytometry results given with Supplementary Data Set 4).

We performed both simple linear regression analyses and rank correlation tests (Spearman Rank and Kendall Tau correlations) based on measured 2C values in R version 3.3.1 (R Core Team 2013) testing the relationships of 1) chromosome number vs. genome size and 2) chromosome number vs. DNA content per chromosome.

### Plant Material and Taxon Sampling for Genome Resequencing

The taxon sampling for the genome resequencing analyses aimed at covering all ploidy levels and the whole distribution range of the genus *Cochlearia*, that way including all different ecotypic variants (e.g. montane versus alpine, or limestone versus siliceous bedrock, salt versus sweet water). We selected a total of 65 samples (Supplementary Data Set 2). *C. islandica* was treated as a separate species, tetraploid populations of British *C. pyrenaica* were referred to as *C. pyrenaica* subsp. *alpina* (abbreviated as *C. alpina* hereafter) and sub-species levels in *C. officinalis* samples from Scandinavia were elided. According to this taxonomic treatment, 19 taxa were included in the study, comprising all accepted species as listed in BrassiBase (Kiefer et al. 2013; see Supplementary Table 1 for a list of accepted species and sub-species) and representing also respective species ranges. Every population/accession is represented by one sample and three samples/taxa from the sister genus *Ionopsidium* were selected to serve as outgroup.

Leaf samples were either selected from own collections (stored as herbarium vouchers, silica-dried material or as living plants) or they were received from other institutions in form of herbarium specimen, silica-dried samples and/or seed material, which was then grown at the Botanical Garden Heidelberg.

A detailed summary of all sequenced samples with information on sample locations and the respective type of leaf material is given with Supplementary Data Set 2.

### DNA extraction and NGS sequencing

The Invisorb Spin Plant Mini Kit (STRATEC Biomedical AG, Birkenfeld, Germany) was used for extractions of DNA from either herbarium, silica-dried or fresh leaf material. Five samples were enriched for chloroplasts prior DNA extraction using a Percoll step-gradient centrifugation (modified from Jansen et al. [2005]).

Library preparation (total genomic DNA) and sequencing were performed at the CellNetworks Deep Sequencing Core Facility (Heidelberg) with library insert sizes of 200 to 400 bp. DNA fragmentation was performed on a Covaris S2 instrument. Either the TruSeq Kit (Illumina Inc., San Diego California, U.S.) or the NEBNext Ultra DNA Library Prep Kit for Illumina (formerly NEBNext DNA Library Prep Kit for Illumina; New England Biolabs Inc., Ipswich, Massachusetts, U.S.) and the NEBNext Multiplex Oligos for Illumina were used for library preparations. Sequencing of multiplexed libraries (six to twelve samples per lane) was performed on an Illumina HiSeq 2000 system in paired-end mode (100 bp).

### NGS data analysis

#### K-mer analysis for ploidy estimation of *C. tridactylites*

We performed a *k*-mer analysis using Jellyfish (Marçais and Kingsford 2011) for all sequenced samples of *C. tridactylites* in order to predict the species-specific genome size. After adapter and quality trimming of raw reads using Trimmomatic version 0.32 (Bolger et al. 2014; LEADING:20, TRAILING:20, SLIDINGWINDOW:4:15, MINLEN:50), the distribution of the *k*-mers 17 and 25 were estimated using Jellyfish. The resulting histograms were analyzed and visualized in R version 3.3.1 (see Supplementary Figure 26).

### Phylogenomic analyses

#### Chloroplast genome assemblies and annotation

A combination of both *de novo* assemblies and reference-based mappings was used for the reconstruction of complete chloroplast genomes of 68 samples. We used the CLC Genomics Workbench version 6.0.4 (CLC Bio, Aarhus, Denmark) for quality and adapter trimming with a quality score limit of 0.001 (corresponding to a phred score of 30) and the minimum read length set to 50 bp. De novo assemblies were performed in CLC for paired trimmed reads (length and similarity fractions set to 0.8). Chloroplast contigs were identified via BLASTn analysis (default settings) and aligned manually using PhyDE v0.9971 (Muller et al. 2010). Gaps between non-overlapping contigs were filled from a complete chloroplast genome that served as a reference, and trimmed reads were mapped back against the so-created preliminary genomes using the CLC *Map Reads to Reference* tool with length and similarity fraction set to 0.9. After manually checking the mapping quality, we performed a CLC variant detection with a variant probability of 0.1 and remaining mapping errors were corrected manually. As an additional quality control, we ran reference-based mappings using the *bwa-mem* algorithm as implemented in BWA version 0.7.8 (Li 2013; default parameters for matching score (1), mismatch penalty (4), gap open penalty (6), gap extension penalty (1) and clipping penalty (5); penalty for an unpaired read pair set to 15). Ambiguously mapped reads and putative PCR duplicated were removed via SAMtools version 0.1.19 (Li et al. 2009; Li 2011) and GATK (McKenna et al. 2010) was applied for a local realignment and an evaluation of per base mapping quality. Finally, consensus sequences were extracted via GATK and masked based on the respective mapping quality using the *maskfasta* tool implemented in BEDtools version 2.19 (Quinlan and Hall 2010; Quinlan 2014).

For the annotation of complete or nearly complete chloroplast genomes, the annotated chloroplast genome of *Arabidopsis thaliana* (NC_000923) served as an initial reference and was therefore aligned to a *Cochlearia* chloroplast genome using MAFFT v7.017 (Katoh et al. 2002; Katoh et al. 2005; Katoh and Standley 2013) as implemented in Geneious version 7.1.7 (Biomatters Ltd., Auckland, New Zealand). We used the FFT-NS-I x1000 algorithm with a 200PAM / k=2 scoring matrix, a gap open penalty of 1.53 and offset value of 0.123 and transferred annotations from the reference in Geneious via the *Transfer Annotation* tool (required similarity 65%). We then checked and adjusted all transferred annotations manually. After the first annotation, the remaining chloroplast samples were aligned to and annotated from the most closely related *Cochlearia* chloroplast genome.

### Chloroplast genome alignments and phylogenetic trees

Sequences of 68 annotated and quality masked cp genomes were aligned in Geneious using MAFFT v7.017 with settings as above. We used Gblocks v0.91b (Castresana 2000) to subject alignment blocks of exons, introns and intergenic spacers to an automatic alignment quality control (minimum block length of 5; gap positions allowed in up to 50% of the samples). Four putative pseudo-genes (trnH, ndhK, rrn16S, rrn23S) were excluded manually as well as three AT-rich regions of poor mapping quality (within trnE-trnT intron, rpl16 intron and trnH-psbA intergenic region). A search for the best partitioning schemes and substitution models for the final 258 blocks was performed using PartitionFinder v1.1.1 (Lanfear et al. 2012; Lanfear et al. 2014) with branch lengths set to be unlinked. We used RAxML version 8.1.16 (Stamatakis 2014) for phylogenetic tree-building (1000 bootstrap replicates) using the GTR+Γ model for the partitioning schemes as determined by PartitionFinder and with three *Ionopsidium* samples set as outgroup.

### Divergence time estimation based on whole chloroplast genome alignment

In the absence of reliable fossil records within the Brassicaceae family (Franzke et al. 2016), a subset of five *Cochlearia* samples and one *Ionopsidium* sample were included in a large-scale divergence time estimation analysis on taxa from the whole Superrosidae clade by Hohmann et al. (2015). The analysis was based on 73 conserved chloroplast genes (51 protein-coding genes, 19 tRNAs, three rRNAs) and allowed for four reliable fossil constrains (also used by e.g. Njuguna et al. 2013; Magallón et al. 2015), to be placed as primary calibration points along the tree (see Hohmann et al. 2015 for details on taxon sampling and data analysis). Namely, minimum ages of calibration points were the *Prunus*/*Malus* split set to 48.5 mya (Benedict et al. 2011), the *Castanea*/*Cucumis* split set to 84 mya (Sims et al. 1999), the *Mangifera*/*Citrus* split set to 65 mya (Knobloch and Mai 1986) and the *Oenothera*/*Eucalyptus* split set to 88.2 mya (Takahashi et al. 1999). The root age of the tree was set to 92-125 mya (uniform distribution) in accordance with Magallón et al. (2015).

Based on this foregoing analysis, we selected secondary calibration points for divergence time estimation within the whole Cochlearieae chloroplast dataset as follows: the *Ionopsidium*/*Cochlearia* split was set to 10.81 mya and the *Cochlearia* crown age was set to 0.71 mya (normal distributions). Divergence time estimation was performed in BEAST 1.7.5 (Drummond et al. 2012) based on the alignment of whole chloroplast genomes (122,798 bp), with partitioning schemes as received from PartitionFinder (see above). Independent site and clock models and a combined partition tree were selected for the two partitions (GTR+Γ) and in order to allow for varying rates among branches, an uncorrelated lognormal relaxed clock model with estimated rates was applied (Drummond et al. 2006). We performed two independent MCMC runs with 100 million generations each (samples taken every 10,000 generations). After combining the two tree files in LogCombiner version 1.5.5 (Drummond et al. 2012) with a burn-in of 50,000,000 generations, treeAnnotator version 1.7.5 (Drummond et al. 2012) was used to combine the 18,000 generated trees to a maximum clade credibility tree which was finally visualized in FigTree version 1.4.1 (Drummond et al. 2012).

### Mitochondrial genome data analysis

Given the challenges in assembling plant mitochondrial genomes *de novo*, mainly caused by the high amount of repetitive regions (Schatz et al. 2010; Straub et al. 2012), we followed an approach by Straub et al. (Straub et al. 2011), which combines the *de novo* assembly of mitochondrial consensus sequences and reference-based mappings to these contigs, serving as a partial mitochondrial reference genome. Therefore, eight long mitochondrial contigs of one *Cochlearia* sample (Cmica_0979; Supplementary Table 12) were retrieved from a *de novo* assembly performed with CLC (same settings as specified above for chloroplast genome *de novo* assemblies) and identified via BLASTn. The CLC *Extract Consensus Sequence* tool was used to extract contigs (minimum coverage threshold of 10x) and annotation of genes and coding sequences was performed with the Mitofy Webserver (Alverson et al. 2010) under default settings. The final reference had a total length of 307,510 bp covering 32 protein-coding genes (some of them partial) and 15 tRNA sequences (13 different tRNAs).

Sequencing reads were trimmed using Trimmomatic (see above) and reference-based mappings were performed with BWA-MEM (mapping settings as specified above for chloroplast genomes). A *Cochlearia* chloroplast genome as well as the nuclear genome of *Arabidopsis thaliana* (NC_003070, NC_003071, NC_003074, NC_003075, NC_003076) were included as additional references in order to reduce mis-mapping of reads originating from pseudogenes. After a mapping quality improvement (same steps as described for chloroplast genome assemblies) the GATK tool *CallableLoci* with a minimum coverage of 20x and a minimum mapping quality of 30 was used to identify regions of high mapping quality. Hereafter, we extracted contig sequences using the GATK tools *UnifiedGenotyper* (output mode EMIT_ALL_SITES) and *FastaAlternateReferenceMaker.* Regions that had not passed the *CallableLoci* quality filters were masked in the final fasta files using BEDtools version 2.19 (Quinlan and Hall 2010; Quinlan 2014) *maskfasta.* Seven *Cochlearia* samples and two *Ionopsidium* outgroup samples with more than 50% missing data were excluded from the mitochondrial phylogenetic analysis.

Geneious was used to concatenate the eight contig alignments and after manually excluding putative pseudogenes, regions of poor alignment quality were removed via Gblocks (minimum block length set to 10). After excluding positions that were masked in all samples, the final alignment had a length of 232,036 bp covering 306 parsimony informative SNPs.

A maximum likelihood analysis was performed with RAxML version 8.1.16 (Stamatakis 2014) with 1000 rapid bootstrap replicates and the GTR+Γ+ I model, selected as the most appropriate substitution model with jModelTest version 2.1.7 (Posada 2008; Darriba et al. 2012).

In order to compare the two organellar phylogenies, we generated a tanglegram using dendroscope version 3.7.2 (Huson and Scornavacca 2012) after collapsing branches with bootstrap support below 95%. The plastid phylogeny was reduced to match the taxon set of the mitochondrial phylogeny using the function ‘drop.tip’ as implemented in the R package ‘ape’ version 5.1 (Paradis and Schliep 2019).

### Nuclear genome data analysis

#### Mapping approach and SNP calling

In order to generate a comprehensive SNP dataset for the nuclear genome, we performed reference-based mappings against the published transcriptome of *Cochlearia pyrenaica* (total length: 58,236,171 bp; Lopez et al. 2017). Therefore, trimmed reads were mapped to the reference using BWA-MEM (Trimmomatic and mapping settings as specified above). Mapping quality was improved and investigated using SAMtools version 0.1.19 and the GATK tools *RealignerTargetCreator*, *IndelRealigner*, *DepthOfCoverage.* For further analyses, the minimum coverage was set to 2x and an upper 2% of coverage cutoff was used for masking high coverage sites for every sample individually, thereby excluding organellar and rDNA transcripts. Three *Cochlearia* samples (Caes_0160, Coff_1289, Ctat_1017) and two *Ionopsidium* samples (Iabu_1074 and Iacau_1072) were excluded from nuclear genome data analysis due to low overall mapping quality (less than 40% of the reference-based mapping positions fulfilling quality and coverage requirements). The *Ionopsidium megalospermum* sample (Imega_1776) also failed the 40% cutoff (∼37% sites callable) but was kept in the analysis in order to have an outgroup sample included.

Callable sites of the respective samples were combined via the *multiIntersectBed* command implemented in BEDtools version 2.19 and SNP callings were performed in sites passing the chosen quality requirements in all samples. GATK’s *UnifiedGenotyper* was used for transcriptome-wide SNP callings in both coding and non-coding regions of the remaining 63 samples. Since ploidy levels were unknown for some of the samples, all samples were treated as diploids. While this approach should not significantly affect the SNP calling in autopolyploids, it is likely to cause some allele drop-outs in allopolyploids. Yet, compared to the general drop-out caused by the low coverage of the sequencing data, this effect will probably be small and preliminary tests using ploidy settings adjusted to respective (known) ploidy levels (data not shown) did not significantly improve the respective SNP callings with regard to the number of called heterozygous sites.

The initial SNP calling revealed 1,250,109 raw SNPs. We first used vcftools to remove variants with a minor allele count less than three, thereby excluding sequencing errors and private SNPs which are undistinguishable in low sequencing depth data. The remaining 492,531 variant sites were filtered using GATKs *VariantFiltration* according to the GATK Best Practices quality recommendations (QD < 2.0 || FS > 60.0 || MQ < 40 || MQRankSum < -12.5 || ReadPosRankSum < -8.0) and keeping only biallelic sites. This resulted in a total of 447,919 hard-filtered variant sites (‘all samples’).

### STRUCTURE analysis

Genetic clustering within the dataset was investigated using STRUCTURE version 2.3.4 (Pritchard et al. 2000; Falush et al. 2003). The analysis was restricted to the 62 *Cochlearia* samples and so *Ionopsidium megalospermum* was excluded from the generated ‘all samples’ vcf file, leaving 400,071 variants to be analyzed. STRUCTURE was run with a burn-in of 5,000 cycles followed by 5,000 iterations per run under an admixture model. We performed ten runs for each *K* from *K* = 1 to *K* = 10 and every subset was analyzed twice in order to test both the *correlated* and the *independent allele frequencies* model. The *correlated allele frequencies* model is supposed to be more sensitive to discrete population structure, yet it might possibly lead to over-estimates of *K* (Pritchard et al. 2010). Therefore, both models were tested and compared. We used the structure-sum script (Ehrich 2006) in R to infer the optimal number of clusters for each analysis according to the Evanno method (Evanno et al. 2005). Results of the different STRUCTURE analyses were processed using the python script structureHarvester.py version 0.6.94 (Earl 2012) and CLUMPP version 1.1 (Jakobsson and Rosenberg 2007) was used to summarize replicate runs of the optimal *K*.

As an example for the evolution of a putatively allopolyploid species within the genus *Cochlearia*, we performed a separate STRUCTURE analysis for *Cochlearia bavarica* and its putative parental species *C. pyrenaica* and *C. officinalis*. We therefore selected the two *C. bavarica* samples as well as three samples of *C. pyrenaica* and *C. officinalis* respectively from the ‘all samples’ vcf file using GATK’s *SelectVariants* tool. After respective SNP filtering steps, 103,874 variant sites remained, and two STRUCTURE analyses (*correlated*/*independent allele frequencies*) were performed with settings as described above for *K* from *K* = 1 to *K* = 6. The optimal *K* was then determined as described above.

### RAxML analysis

To further analyze phylogenetic relationships based on the nuclear genomic data, we performed a ML tree reconstruction with RAxML version 8.1.16 (Stamatakis 2014). In order to run RAxML with an ascertainment bias correction, ambiguous sites had to be removed from the ‘all variants’ vcf file, resulting in 298,978 remaining variant sites. JModelTest version 2.1.10 was utilized to determine the best-fit nucleotide substitution model. The RAxML analysis was carried out under the GTR+Γ substitution model with 1,000 rapid bootstrap replicates, and FigTree version 1.4.1 (Drummond et al. 2012) was used for visualization of the best final ML tree.

### SplitsTree analysis

Aside from the ML tree search, we used SplitsTree version 4.15.1 (Huson 1998; Huson and Bryant 2006) to investigate conflicting or reticulate phylogenetic relationships. The input file was generated from the ‘all samples’ vcf file (447,919 hard-filtered variant sites) using the Python script ‘vcf2phylip’ (Ortiz 2019). We used the NeighborNet algorithm based on uncorrected p-distances and equal angles to compute the split network.

### TreeMix analysis

Moreover, we used TreeMix version 1.13 (Pickrell and Pritchard 2012) to further investigate the historic relationships among *Ionopsidium* and the analyzed *Cochlearia* accessions. The software is designed for the estimation of population trees with admixture, yet it can also be used with only a single individual representing a population. We therefore turned off the correction for sample size (-noss) as this could lead to an overcorrection with only one sample per population. The analysis was based on 447,919 hard-filtered variant sites (see above) and we allowed for 0 to 10 migration edges (m). *Ionopsidium* was set as root and we used a SNP window size of 100 (-k) for all analyses. 10 initial runs were performed for every m and the R package ‘optM’ (Fitak 2019) was used to identify the optimal number of migration events based on the Evanno method (Evanno et al. 2005). For the best m, we then performed 100 TreeMix runs as bootstrap replicates and a consensus tree was inferred from the generated 100 maximum likelihood trees using the program SumTrees version 4.10 (Sukumaran and Holder 2010).

### ABC modelling - demographic history of diploid arctic and central European *Cochlearia*

The genetic history of diploid lineages of *Cochlearia* from the arctic (ARC) and central European (EUR) distribution ranges was analyzed in a coalescent modelling framework with Approximate Bayesian Computation (ABC; Tavaré et al. 1997; Beaumont et al. 2002). We first evaluated a set of four models differing in the history of gene flow (see Figure 3c), and in a second step fitted parameters for relevant models.

Observed data were prepared by filtering all individuals for read coverage with a minimum coverage of 4x and an upper coverage cutoff of 2%. In order to restrict the data to sites that follow a model of neutral evolution as close as possible, only silent (four-fold degenerate) sites were retained. These included bi-allelic SNPs as well as monomorphic sites, to characterise neutral genetic variation in an absolute sense. Contigs with any SNPs displaying excessive heterozygosity (Hardy-Weinberg test; Danecek et al. 2011) indicative of paralogs were excluded. This resulted in a dataset with 22 diploid EUR individuals, 12 diploid ARC individuals, and 6,387 sites located in 5,601 independent contigs with a minimum length of one base pair. Summary statistics for the observed data were calculated with the same code as used for simulations.

Coalescent simulations were carried out in a custom pipeline based on the work by (Roux et al. 2013), using the simulator msnsam (Hudson 2002; Ross-Ibarra et al. 2008). We used a set of 114 population genetic summary statistics, among them a folded two-dimensional site frequency spectrum (Gutenkunst et al. 2009).

The sampling scheme of observed data, i.e. number of samples, contigs and their lengths, was replicated identically in the simulations. Additional fixed parameters were the mutation and recombination rates, which were both set to 6.51548*10^-9 per site per generation, the silent site mutation rate for Brassicaceae (De La Torre et al. 2017). Although we truncated the original contigs to their silent sites only, recombination rates (but not mutation rates) were specified such that they reflected the original contig lengths with all sites. Free parameters of the four models were sampled from uniform prior distributions and are listed in Supplementary Table 13. All of these models put the history of effective population sizes of ARC and EUR populations in two phases, one ancient phase during which Ne was constant, and one recent phase in which Ne either remained the same as before, or exponentially grew or exponentially declined towards the present.

To evaluate the observed data against four alternative models, we used the R package ‘abcrf’ (Pudlo et al. 2016) with 40,000 simulations per model and 1,000 trees per random forest, subsampling 100,000. Whereas traditional rejection ABC model choice approximates a models posterior probability based on the relative frequency of simulations under the model that are similar to the observed data, the random forest ABC (ABC-RF) posterior probability is an estimate of the classification error, i.e. the probability that the resulting classification is correct. Model parameters were also estimated using ‘abcrf’ with 100,00 simulations and separate regression random forests (500 trees) for each parameter. Goodness of fit was evaluated by posterior predictive checks, with 40,000 new simulations generated from the full approximated posterior distributions, and the standardized Euclidean distances between observed data and posterior simulations as done by the rejection-ABC function from the R package ‘abc’ (Csilléry et al. 2012).

### GC-MS primary metabolite profiling

#### Plant Material and Taxon Sampling for Metabolite Profiling

For the GC-MS-based metabolite profiling, we selected 14 *Cochlearia* taxa from a total of 28 populations/accessions representative of the total ecotypic variation within the genus (Supplementary Table 9). Additionally, 5 populations/species of the genus *Ionopsidium* were included as an outgroup. Whenever possible, at least 4 plants per accession were analyzed but, in few cases, smaller sample sizes were accepted due to limited seeds/living plant material. A total of 141 plants, either collected in the wild or grown from seed material was considered for metabolomic analyses. Prior to the experiments, all plants were cultivated under greenhouse conditions in a substrate consisting of seedling potting soil, quartz sand, and either composted earth or a peat-based substrate (Ökohum GmbH, Herbertingen, Germany).

### Habitat characterization and ecotype definition

For all 28 *Cochlearia* populations, bioclimatic variables were downloaded from the high-resolution climate data WorldClim grids (http://www.worldclim.org; Hijmans et al. 2005) at a resolution of 30 arcseconds (∼1 km^2^ / pixel). Since geographical coordinates were not available for most of the *Ionopsidium* accessions included in the metabolite profiling, we excluded *Ionopsidium* from the habitat characterization and instead treated it as a Mediterranean ecotype in downstream analyses. For *Cochlearia*, nine out of 19 standard topo-climatic variables were selected for hierarchical cluster analysis in order to assign the populations to common climatic ecotypes (Source Data). These include the temperature-related bioclimatic variables Annual Mean Temperature (BIO1), Max Temperature of Warmest Month (BIO5), Min Temperature of Coldest Month (BIO6), Mean Temperature of Warmest Quarter (BIO10) and Mean Temperature of Coldest Quarter (BIO11) as well as four temperature/precipitation-related variables, namely Mean Temperature of Wettest Quarter (BIO8), Mean Temperature of Driest Quarter (BIO9), Precipitation of Warmest Quarter (BIO18), Precipitation of Coldest Quarter (BIO19).

A hierarchical cluster analysis based on the selected population-specific bioclimatic variables was performed in R version 3.3.1. Prior to scaling and log transformation, a constant value (+150) was added to the original variables in order to avoid negative values. The dist() function was used to compute euclidian distances and a hierarchical clustering was performed using hclust() (‘stats’ package) according to the Ward method (“ward.D2”; Ward Jr 1963; Murtagh and Legendre 2014) which has been applied for climate data cluster analyses before (Unal et al. 2003). The best number of clusters in the dataset was evaluated using the package ‘NbClust’ (Charrad et al. 2014) with all 26 indices being computed, and the final cluster dendrogram was visualized using the R package ‘factoextra’ (Kassambara 2015).

### Temperature treatment

The metabolomic experiments were performed in two rounds (first batch: 27 *Cochlearia* accessions; second batch: 1 *Cochlearia* accession, 5 *Ionopsidium* accessions) with identical experimental setup (Figure 4c). After an initial phase of acclimatization for two weeks in a plant room with a 16/8 h day/night, 20°C/18°C day/night cycle, leaf samples were harvested from all 141 plants, frozen in liquid nitrogen and transferred to -80°C until metabolite extraction. Hereafter, a cold treatment was performed on half of the plants of each accession in a cold chamber with a 16/8 h day/night, 5°C day/night cycle, while the remaining plants stayed under control conditions. Leaves from all plants (cold and control) were collected again after 20 days and treated as described above.

### GC-MS-based metabolite profiling

Metabolite profiling was performed using gas chromatography-mass spectrometry (GC-MS) and primary metabolite extraction and analysis steps as described by Roessner et al. (2001). Briefly, 15 – 40 mg of the previously collected and frozen leaf material were homogenized by grinding in liquid nitrogen and hereafter mixed with 360 µL cold methanol. 20 µg of ribitol were added as an internal normalizing standard. After extracting the sample for 15 min at 70°C, it was mixed thoroughly with 200 µL chloroform and 400 µL water and centrifuged subsequently. 200 µL of the methanol-water upper phase containing polar to semi-polar metabolites were collected and concentrated to dryness in a vacuum concentrator. A two-step derivatization procedure including methoximation of the dried residue followed by silylation was performed (Lisec et al. 2006). To this end, the residue was first re-suspended in a methoxyamine-hydrochloride/pyridine solution for a methoxymization of the carbonyl groups. The sample was then heated for 90 min at 37°C and further silylated with N-methyl-N-trimethylsilyltrifloracetamide at 37°C for 30 min.

GC-MS analysis was performed on a gas chromatograph system equipped with quadrupole mass spectrometer (GC-MS-QP2010, Shimadzu, Duisburg, Germany). For this, 1 µL of each sample was injected in split mode with a split ratio of 1:20 and the separation of derivatized metabolites was carried out on a RTX-5MS column (Restek Corporation, Bellefonte, Pennsylvania, USA). Metabolites were detected using optimized instrumental settings (Lisec et al. 2006).

### GC-MS data processing

A two-pronged approach was employed for metabolite annotation. Briefly, obtained raw data files were first converted into an ANDI-MS universal file format for spectrum deconvolution and compound identification via the reference collection of the Golm Metabolome Database (GMD, http://gmd.mpimp-golm.mpg.de/) using the AMDIS program (Automated Mass Spectral Deconvolution and Identification System; www.amdis.net/). Kovats retention indices were calculated for deconvoluted mass spectra from measurements of an alkane mixture and hereafter compared with best hits obtained via the GMD database. The Shimadzu GCMS solutions software (v2.72) interface was further used for manual curation of metabolite annotation versus an in-house library of authentic standards analyzed under the above analytical conditions.

CSV output files were exported for each measurement batch with peak areas obtained for quantifier ions selected for a set of 40 compounds (annotated as known compounds by the above annotation approach or considered as unknown compounds) consistently detected in all analyzed samples. Peak areas (Supplementary Data Set 3) were scaled on a sample-basis according to the extracted amount of leaf tissue and further in percent of the peak area of the ribitol internal standard, the latter to account for putative extraction and analytical performance variations across the different measurement batches. Cross-sample variations of the ribitol peak area did not differ significantly between the different measurement batches (between-batch *F* = 1.998, one-way ANOVA *P* = 0.1586) and relative standard deviation of this internal standard did not differ more than 15% between measurement batches. While not providing absolute quantification information on individual compounds, the normalized compound table allows for cross-condition statistical analysis of metabolite relative changes. To this end, normalized tables were concatenated in one matrix prior to subsequent univariate and multivariate statistics (Supplementary Data Set 3).

### Multivariate statistical analysis of metabolite data and integration of bioclimatic population clusters

The aov() function in R was used to perform separate one-way ANOVAs on the normalized metabolite data grouped by the four bioclimatic population clusters and *Ionopsidium* to investigate differences in the compound concentrations between control and cold treatment in the different clusters. To further investigate putative metabolomic differentiation between the clusters, PCAs were carried out on the normalized metabolite data using the dudi.pca() function implemented in R package ‘ade4’ (Dray and Dufour 2007). We then performed DAPCs (Jombart et al. 2010) via the function dapc() embedded in the R package ‘adegenet’ (Jombart 2008; Jombart and Ahmed 2011). Input data was centered and scaled and prior to each DAPC a cross-validation was performed using the xvalDapc() function of the ‘adegenet’ package with 1000 replicates in order to find the optimal number of PCs to retain. The latter was determined from the lowest root mean squared error associated with the predictive success. Group priors were first defined by temperature treatment in order to analyze the general response to the different conditions. For further DAPCs, group priors were set to represent the four bioclimatic clusters and *Ionopsidium*. DAPCs were performed separately for compound data from the second measurement of either control plants (20°C) or cold treated plants (4°C). All discriminant functions were retained and DAPC results were visualized using the R function scatter() from the ‘ade4’ package.

## Data availability

Raw sequencing data generated for the current study is available under ENA/GenBank bioproject PRJEB21320 [https://www.ncbi.nlm.nih.gov/bioproject/PRJEB21320], and annotated plastome sequences under ENA/GenBank accession numbers LT629868 - LT629930 and LN866844 - LN866848 (see Supplementary Data Set 2). All Supplementary Data Sets (1-13) including the annotated mitochondrial consensus sequences as well as metabolite profiling data and input files for NGS data analyses are available at Dryad [doi:10.5061/dryad.fbg79cnsn]. FCS files of flow cytometry data are available at FlowRepository under identifier FR-FCM-Z3FY.

Data supporting the findings of this study are contained within the paper and its Supplementary Information file. The data underlying the main and Supplementary Figures are provided as Source Data.

## Acknowledgements

We thank our gardeners Thorsten Jakob, Bärbel Schwarz and Frank Korn for excellent plant curation. Peter Sack, Lisa Kretz, Florian Michling and Lua Lopez are acknowledged for assistance as is David Ibberson at Heidelberg Deep Sequencing Core facility and Rainer Schulz for help with cytogenetic compilations. This work was supported by the German Research foundation (DFG; KO2302/13 and KO2302/16 to M.A.K.).

## Author contributions

M.A.K. conceived and designed the study. E.M.W., E.G., M.S. and M.A.K. performed experiments and collected data. E.M.W., E.G., M.S., L.Y. and M.A.K. analyzed data or contributed with conceptual ideas. E.M.W., M.A.K. and M.S. wrote the manuscript. All authors contributed to the final version of the manuscript.

## Materials & Correspondence

Correspondence and material requests should be addressed to M.A.K.

